# Shorter unreported sequences in a RACE-Seq study involving seven tissues confirms ∼150 novel transcripts identified in MCF-7 cell line PacBio transcriptome, leaving ∼100 non-redundant transcripts exclusive to the cancer cell line

**DOI:** 10.1101/104257

## Abstract

PacBio sequencing generates much longer reads compared to second-generation sequencing technologies, with a trade-off of lower throughput, higher error rate and more cost per base. The PacBio transcriptome of the breast cancer cell line MCF-7 was found to have ∼300 transcripts un-annotated in the current GENCODE (v25) or RefSeq, and missing in the liver, heart and brain PacBio transcriptomes [1]. RACE-sequencing (RACE-seq [2]) extends a well-established method of characterizing cDNA molecules generated by rapid amplification of cDNA ends (RACE [3]) using high-throughput sequencing technologies, reducing costs compared to PacBio. Here, shorter fragments of ∼150 transcripts were found to be present in seven tissues analyzed in a recent RACE-seq study (Accid:ERP012249) [4]. These transcripts were not among the ∼2500 novel transcripts reported in that study, tested separately here using the genomic coordinates provided, although ‘all curated novel isoforms were incorporated into the human GENCODE set (v22)’ in that study. Non-redundancy analysis of the exclusive transcripts identified one transcript mapping to Chr1 with seven different splice variants, and erroneously mapped to Chr15 (PAC clone 15q11-q13) from the Prader-Willi/Angelman Syndrome region (Accid:AC004137.1). Finally, there are ∼100 non-redundant transcripts missing in the seven tissues, in addition to other three tissues analyzed previously. Their absence in GENCODE and RefSeq databases rule them out as commonly transcribed regions, further increasing their likelihood as biomarkers.

## Introduction

The obvious advantages of long-read sequencing technologies [5] is currently tempered by the costs, low throughput and possible higher error rates [6]. Pacific Biosciences (PacBio) sequencing [7] generates much longer reads compared to second-generation sequencing technologies [8], with a trade-off of lower throughput, higher error rate and more cost per base [6, 9]. The longer sequence lengths in PacBio compared to other sequencing methods might alleviate assembly issues associated with other methods with shorter read lengths [10, 11]. Another innovative technique extends a well-established method for characterizing cDNA molecules generated by rapid amplification of cDNA ends (RACE [3]) using high-throughput sequencing technologies to reduce time and cost, and increase sensitivity [2, 12, 13].

The annotation of the human genome is an ongoing collaborative effort - two major independent annotation databases, periodically updated, are RefSeq [14] and GENCODE [15]. Genome annotation is critical for correlating disease to genomic variants [16], since transcribed regions, aside from coding for proteins, play significant regulatory roles in the cellular machinery [17]. Other transcriptional mechanisms like alternate splicing play an important role in cellular regulation [18]. Dysregultion and aberrations in splicing is implicated in cancer [19–22].

The fast reducing costs of genome sequencing has resulted in the availability of unprecedented volumes of data, necessitating the development of different pipelines to process and analyze this data. The under-utilization of transcriptomes while annotating genomes [23–25] was recently emphasized for the walnut genome [26]. PacBio has provided open access to the transcriptome of the MCF-7 breast cancer cell line [27, 28]. There are currently two versions (2013 and 2015). Previously, the 2013 version of the the MCF-7 transcriptome was used to find *∼*300 transcripts that have no annotation in the current RefSeq and GENCODE databases, and predominantly absent in heart, liver and brain transcriptomes also provided by PacBio [1]. Also, *∼*200 transcripts were not present in a recent catalogue of un-annotated long non-coding RNAs (lncRNA) from 6,503 samples (*∼*43 Terabases of sequence data) [29]. Recently, using known long non-coding RNA (lncRNA) loci, RACE fragments were sequenced using Roche 454 FLX+ platform to annotate *∼*2500 novel lncRNAs after analyzing seven tissues (brain, heart, kidney, liver, lung, spleen and testis) [4]. There were no common transcripts in this study among the *∼*300 transcripts as determined using the genomic coordinates provided for these novel *∼*2500 transcripts, as expected since they were included in the GENCODE v22.

Here, the sequencing data from seven tissues (ENA accid:ERP012249) made available by the RACE-seq study [4] were analyzed to compare the *∼*300 un-annotated transcripts detected by YEATS [1]. Shorter fragments of *∼*150 transcripts were found to be present in several of these tissues, but were not among the *∼*2500 novel transcripts reported in that study. Finally, there are *∼*100 non-redundant transcripts missing in the seven tissues, in addition to other three tissues (liver, heart and brain) analyzed previously. Their absence in GENCODE and RefSeq databases rule these out as commonly transcribed regions, further increasing their likelihood as biomarkers.

## Results and discussion

### Common shorter transcripts found in the RACE-seq sequencing data

There are 155 transcripts from the *∼*300 novel transcripts from the MCF-7 cell line, reported earlier [1], which have shorter versions in the RACE-seq study (FILE:common.transcripts.txt, Table 1). However, these were not reported among the *∼*2500 novel transcripts reported earlier. The genomic coordinates are obtained from https://public_docs.crg.es/rguigo/Papers/2016_lagarde-uszczynska_RACE-Seq/ (also provided here in FILE:phase6-clean.bed, n=2486). Some transcripts have homologs in multiple tissues - in such cases, the longest transcript has been used as the representative (Table 1).

**Table 1:** Shorter length transcripts present in the RACE-seq study compared to PacBio transcriptome: RACE-seq ids are based on European Nucleotide Archive Accid:ERP012249. For transcripts that have homologs in multiple tissues, the longest transcript is chosen and the corresponding tissue (LTiss) is mentioned. The complete list of 155 transcripts is in FILE:common.transcripts.txt. NTiss:Number of tissues that the transcript occurs in, PLen: Length of PacBio transcript, RLen: Length of RACE-seq transcript, %I: Percentage identity, LTiss: Tissue with the longest transcript, BBS: BLAST bitscore.

| PacBio id | RACE-seq id | NTiss | PLen | RLen | %I | LTiss | BBS |
| --- | --- | --- | --- | --- | --- | --- | --- |
| CHR1.8078444_8080132+.I1C_C83903.F2P8.1695.RT | ERR1033123.145501 | 7 | 1689 | 968 | 99 | liver | 1722 |
| CHR6.151722733_151724445+.I1B_C47905.F3P12.1470.RT | ERR1033132.186957 | 7 | 1464 | 984 | 99 | spleen | 1742 |
| CHR12.46133248_46135312_.I2A_C109550.F3P18.2072.RT | ERR1033145.416197 | 6 | 2067 | 1021 | 99 | testis | 1844 |
| CHR14.58747258_58748777+.I1C_C41814.F4P18.1523.RT | ERR1033130.210294 | 6 | 1517 | 931 | 99 | spleen | 1640 |
| CHR15.99296643_99298133_.I1C_C49931.F1P41.1499.RT | ERR1033132.279152 | 6 | 1492 | 970 | 99 | spleen | 1775 |
| CHR15.99321470_99323379+.I1C_C77550.F1P14.1914.RT | ERR1033122.20588 | 6 | 1908 | 918 | 99 | kidney | 1639 |
| CHR16.29592837_29594151_.I1C_C44797.F2P9.1319.RT | ERR1033124.128471 | 6 | 1313 | 870 | 99 | lung | 1578 |
| CHR17.61279795_61281644+.I1C_C72580.F1P16.1855.RT | ERR1033132.122017 | 6 | 1848 | 974 | 99 | spleen | 1762 |
| CHR2.32796474_32798035_.I1C_C47523.F2P11.1568.RT | ERR1033132.502245 | 6 | 1562 | 966 | 99 | spleen | 1681 |
| CHR21.34129622_34130794_.I1A_C59236.F1P11.1180.RT | ERR1033147.381934 | 6 | 1175 | 967 | 99 | testis | 1755 |

### Non-redundant set of transcripts exclusive to the breast cancer cell line - alternate splicing

This left *∼*130 (FILE:notAnno.txt) transcripts missing in the liver, heart and brain PacBio transcriptomes [1], and also in the transcriptome of seven tissues sequenced in the RACE-seq study [4]. These transcripts have some redundancy, and include several alternately spliced variants. To identify these, and make the set non-redundant, a grouping algorithm (see Methods) was applied, reducing the number of non-redundant transcripts exclusive to the MCF-7 breast cancer cell line to 107 (FILE:notAnno.nonredundant.list). 37 transcripts have open reading frames *>*100 amino acids (FILE:notAnno.nonredundant.list.orf100.info), and are candidates for protein coding genes.

### Future work

Future work will compare the PacBio transcriptome analyzed here with the RNA-seq derived transcriptome of MCF-7 cells (SRA:SRX701874), part of the study for profiling long noncoding RNAs with targeted RNA sequencing (BioProject:PRJNA261251) [30]. Also, a stand-alone analysis of the RACE-seq data (ENA accid:ERP012249) [4] will be done to identify additional un-annotated transcripts.

### Conclusion

The MCF-7 cell line was derived from pleural effusion (a condition in which excess fluid builds around the lung) of a patient with metastatic mammary carcinoma [27]. The presence of transcripts exclusive to this cell line can be progressively tested to other transcriptomes, obtained using different sequencing methods, to narrow down a possible set of biomarkers. The absence of *∼*100 transcripts in GENCODE and RefSeq databases, in addition to 10 additional transcriptomes, rule these out as commonly transcribed regions, increasing their likelihood as cancer biomarkers. Several of these transcripts are putative protein coding genes.

## Materials and methods

The GENCODE, RefSeq and PacBio datasets were described previously [1]. It is briefly described here again.

### GENCODE/RefSeq/PacBio datasets described previously

GENCODE release 25 was downloaded from https://www.gencodegenes.org/ (release date 07/2016). The RefSeq database was created from https://www.ncbi.nlm.nih.gov/nuccore choosing mRNA, rRNA, cRNA, tRNA and ncRNA sequences (FILE:mrna.refseq.160k.fa, n=161k, REFSEQ. NTDB). The MCF-7 transcriptome was obtained from http://www.pacb.com/blog/data-release-human-mcf-7-transcriptome (2013 version). The PacBio dataset for human heart, liver and brain transcriptomes is available at http://datasets.pacb.com.s3.amazonaws.com/2014/Iso-seq_Human_Tissues/list.html. The transcripts have been renamed to allow Unix style filenames.

### RACE-seq dataset

Recently, RACE-seq, an experimental workflow based on RACE (Rapid Amplification of cDNA Ends) [3] and long read RNA sequencing was used to target 398 distinct long non-coding RNA loci from the GENCODE v7 annotation, on a set of cDNA libraries from seven human tissues (brain, heart, kidney, liver, lung, spleen and testis) [4]. The sequence data was made available at the European Nucleotide Archive (Accid:ERP012249). The curated novel isoforms have been incorporated into the human GENCODE v22 onwards (the current study uses GENCODE v25). Other data was made available at http://public-docs.crg.es/rguigo/Papers/2016_lagarde-uszczynska_RACE-Seq/. Here, four data files from each tissue were combined to created a single BLAST database for each tissue, in order to allow a tissue specific analysis. The YeATS suite was used extensively to query these databases using the BLAST command-line interface [31] with the *∼* 300 novel transcripts identified previously [1].

### Additional algorithms included in the YEATS suite

A grouping algorithm (YeATS-GROUP) was added to the YeATS suite [23]. For a given set of sequences, a BLAST database is created [31]. Each sequence is BLAST’ed to this database, and is linked to another sequence if the BLAST bitscore (BBS) value is more than the specified cutoff (500 in this case). Finally, a group is created such that any sequence in the group has at least one sequence with which it has a homology *>*BBS=500. The BLAST bitscore was used as a comparison metric instead of the Evalue since it allows differentiation for high homologies where Evalue goes to zero.

### External programs

Multiple sequence alignment was done using MAFFT (v7.123b) [32], and figures generated using the ENDscript server [33]. Open reading frames were obtained using the ‘getorf’ program from the EMBOSS suite [34]. Hardware requirements are very modest - all results here are from a simple workstation (8GB ram) and run-times were a few hours.

**Figure 1:**
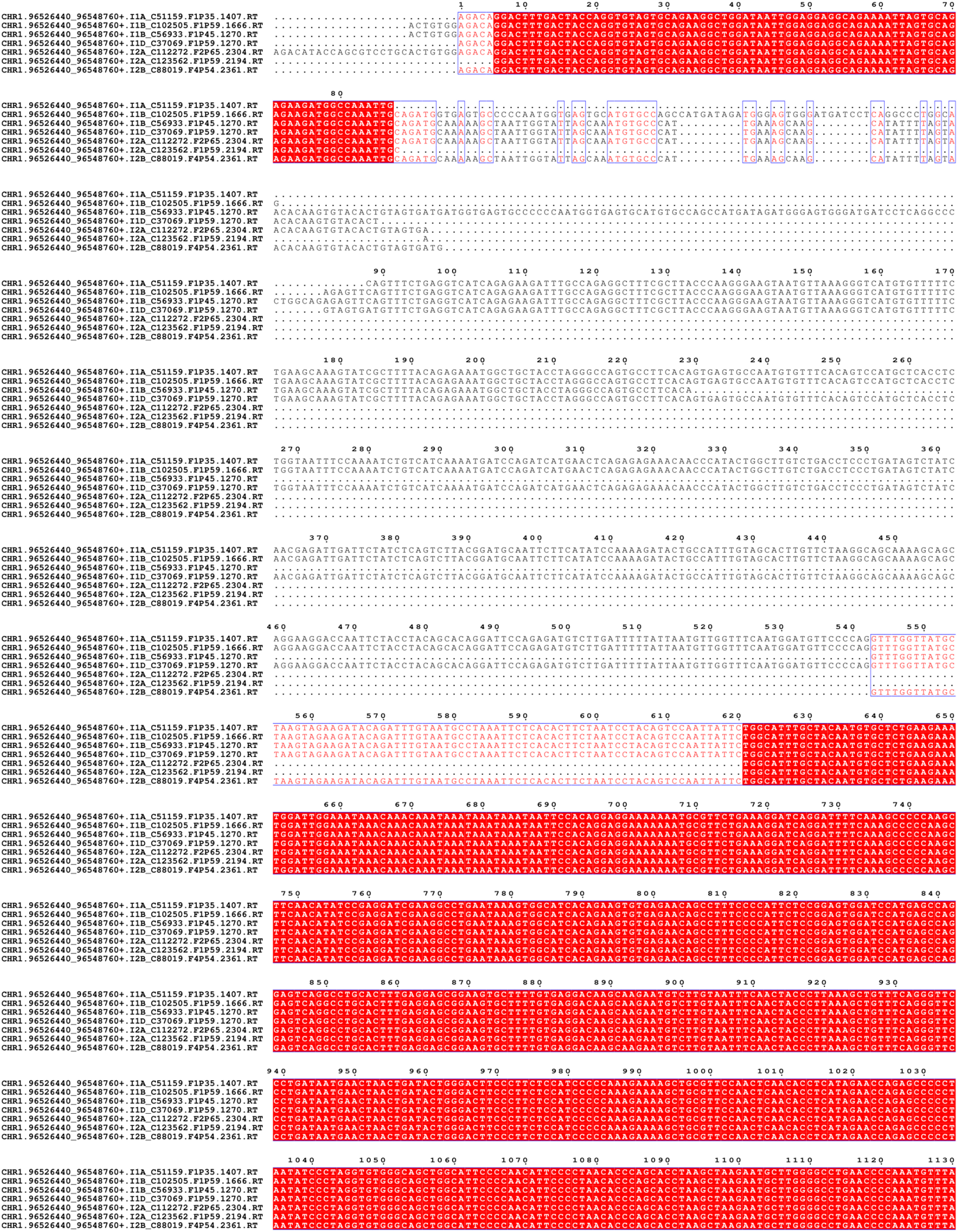
Seven splice variants for one transcipt found in MCF cell lines exclusively: The transcript is absent in liver, heart and brain PacBio transcriptomes, and also seven tissues sequenced in the RACE-seq study. The multiple sequence alignment is truncated, see FILE:MSA.splice.pdf for the full alignment.

